# Optimal transport for mapping senescent cells in spatial transcriptomics

**DOI:** 10.1101/2023.08.16.553591

**Authors:** Nam D. Nguyen, Lorena Rosas, Timur Khaliullin, Peiran Jiang, Euxhen Hasanaj, Jose A. Ovando, Marta Bueno, Melanie Konigshoff, Oliver Eickelberg, Mauricio Rojas, Ana L. Mora, Jose Lugo-Martinez, Ziv Bar-Joseph

## Abstract

Spatial transcriptomics (ST) provides a unique opportunity to study cellular organization and cell-cell interactions at the molecular level. However, due to the low resolution of the sequencing data additional information is required to utilize this technology, especially for cases where only a few cells are present for important cell types. To enable the use of ST to study senescence we developed scDOT, which combines ST and single cell RNA-Sequencing (scRNA-Seq) to improve the ability to reconstruct single cell resolved spatial maps. scDOT integrates optimal transport and expression deconvolution to learn non-linear couplings between cells and spots and to infer cell placements. Application of scDOT to existing and new lung ST data improves on prior methods and allows the identification of the spatial organization of senescent cells, the identification of their neighboring cells and the identification of novel genes involved in cell-cell interactions that may be driving senescence.

## 1 Introduction

Recent advancements in genomics technologies have facilitated the profiling of gene expression at the single-cell level, unveiling valuable insights regarding the molecular heterogeneity of complex biological systems. While single-cell RNA sequencing (scRNA-seq) has significantly enhanced our comprehension of cell-type diversity, it lacks spatial information due to the dissociation of cells. Spatial transcriptomics (ST) techniques enable the preservation of spatial information within tissue samples but typically offer lower resolution or coverage compared to scRNA-seq data. Hence, the integration of scRNA-seq and ST data becomes imperative for acquiring a spatially informed single-cell resolution dataset [28]. This integration approach not only ensures a more comprehensive understanding of the molecular heterogeneity within complex biological systems but also retains the spatial context of gene expression.

Existing methods for integrating single-cell and spatial transcriptomics data primarily focus on cell-type deconvolution. These methods decompose gene expression in a spatial spot into linear combinations of fractions attributed to different cell types, utilizing the single-cell data solely as a reference [24, 12, 30, 21, 5, 29, 2, 10]. While successful, these methods often struggle when it comes to cell types with only a few cells [6, 32, 51]. Moreover, in cases where these smaller cell types are very similar to cell types with larger number of cells, the assignment of deconvolution methods often completely ignore these smaller cell types as shown in Results.

Cellular senescence, a state of permanent growth arrest, is implicated in various age-related diseases. Understanding cellular senescence requires analyzing cell-cell communications at the individual cell level, as the process exhibits heterogeneity, where only a few cells within a given cell type enter a senescent state simultaneously. Additionally, paracrine senescence, in which a senescent cell can induce senescence in neighboring cells, is of significant importance. Effective communication between senescent cells and neighboring cells is crucial for the progression and maintenance of the senescent phenotype [38, 13]. Senescent cells actively engage in intercellular communication, primarily through the secretion of senescence-associated secretory phenotype (SASP) factors, influencing neighboring and distant cells [13, 15]. However, the mechanisms underlying these communications remain poorly understood. To address this gap, and to enable the study of cell-cell interactions for these small number of senescent cells within a cell type using spatial transcriptomics, we propose an innovative computational framework that integrates single-cell and spatial transcriptomics data. This approach allows us to infer cell-cell communications based on the proximity of cells, whether short- or long-range, shedding light on the intricacies of senescence-associated intercellular signaling. This method offers a superior alternative to organoids, where only cell types interact in an artificial environment.

Mapping individual cells to their spatial origins requires fine-grained mapping, which is prone to imprecise results due to the similarity within cell types and the non-linear relationship between gene expression levels in scRNA-seq and spatial transcriptomics [46]. Methods proposed for this task compute a similarity score in a shared latent space. This similarity score is then coupled with a statistical test to determine the significance of the assignment [46, 19]. Other techniques, e.g., canonical correlation analysis or non-negative matrix factorization, for constructing shared latent space have also been used [4, 43, 49]. In contrast, here, we utilize optimal transport [40, 45], a mathematical framework that allows for the comparison and matching of probability distributions. Specifically, we use optimal transport to learn the non-linear coupling between cells and spots by aligning the distributions of gene expression profiles across these two datasets. Our approach employs a probabilistic mapping, where the precision of the mapping is modulated by incorporating the coarse-grained mapping of cell types obtained from the deconvolution task. We solve these two complementary optimization tasks using a bilevel optimization approach [7], based on the differentiable deep declarative network [16] (Figure 1).

**Figure 1:**
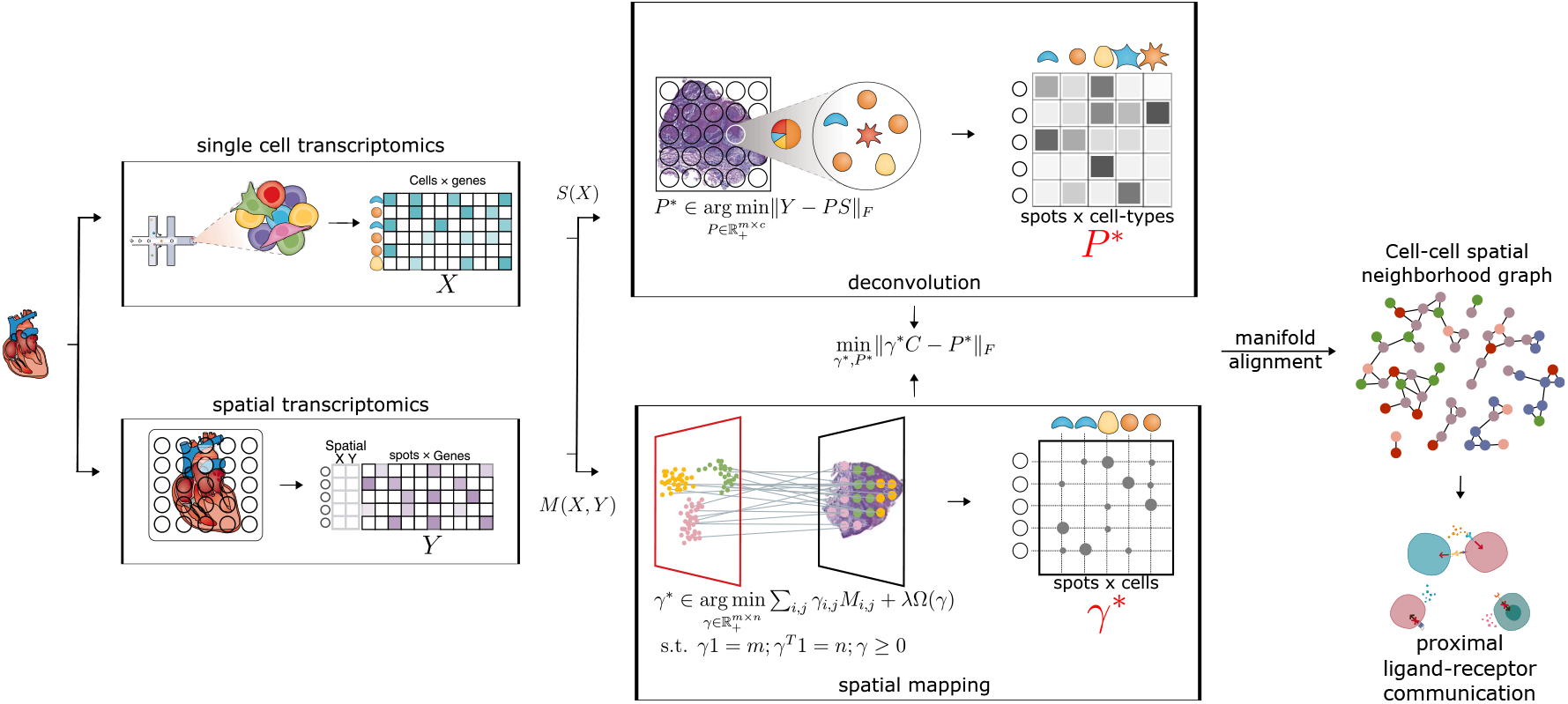
Method workflow: scDOT takes gene expression profiles from a scRNA-seq dataset and a spatial transcriptomics dataset as inputs. Additionally, cell type information for cells in the scRNA-seq data and spatial coordinates for spots in the spatial transcriptomics data are provided. scDOT simultaneously and in parallel learns the cell type fraction of each spot (deconvolution task) and the mapping between individual cells in the scRNA-seq data and individual spots in the spatial transcriptomics data (spatial reconstruction task). The resulting mapping matrix between cells and spots is then utilized to construct the cell-cell spatial neighborhood graph, where cells are connected if they are in close physical proximity.

Our approach incorporates two types of data, namely scRNA-seq and spatial transcriptomics, as inputs. It employs iterative computations to perform cell type deconvolution and cell-to-spot spatial mapping. As a result, it produces a coupling matrix between cells and spots that serves as an initial integration outcome. This coupling matrix is subsequently used to infer the cell-to-cell spatial neighborhood graph by aligning cells with spots possessing known spatial coordinates (see Figure 1). Essentially, the spot coordinates play a crucial role in determining the physical closeness between cells.

We tested scDOT on both, simulated and new spatial data. As we show, it can accurately assign cells to their spot of origin outperforming prior methods for this task. For the new samples for idiopathic pulmonary fibrosis (IPF), scDOT identifies the spatial distribution and cell-cell interactions between senescence and non-senescence cells and the set of genes involved in these interactions.

## 2 Results

We developed an optimal transport (OT) method for mapping scRNA-Seq data to spatial trancriptomics data. The method, illustrated in Figure 1 performs iterative computations for cell type deconvolution and cell-to-spot spatial mapping, resulting in the generation of the coupling matrix *γ* as an upstream integration outcome. This coupling matrix is then utilized to infer the cell-to-cell spatial neighborhood graph by aligning cells to spots with known spatial coordinates.

### 2.1 scDOT efficiently reconstructs individual cells to their spatial origins

We first tested scDOT on two simulation datasets where ground truth is known (Methods). The outcome of reconstructing single-cell data, i.e., the coupling matrix *γ*, when using simulation dataset 1 reveals that it successfully recovers the spatial origins of a high fraction of cells (56% to 76%, depending on a predefined threshold to determine high probability). *γ* represents probabilistic couplings and so a specific cell can be mapped to several location with different probabilities (which sum up to 1). We found that in most cases the distribution *γ*_:,*j*_ exhibits is exteremly heavy-tailed and places a disproportionately high amount of probability densities at 0. We therefore defined a high probability of associating with a location based on distribution properties (99th-, 95th-, 90th-quantile, or the 75th quantile (the third quantile) plus 1.5 times the interquartile range (IQR) (Turkey’s fences)). Obviously, stricter the threshold, the fewer cells that are correctly matched. However, even for a very high cutoff we find very large percentage of correct matches (70% of cells at a threshold above the 90th quantile and 56% of cells at a threshold above the 99th quantile when using synthetic data 1). However, the slower decay of reconstruction results due to a more strict threshold is desirable and can be achieved through a heavier tail in the distribution *γ*_:,*j*_.

In addition, previous studies show that cell type deconvolution methods tend to miss rare cell type [6]. In contrast when using OT we are able to map rare cell types to their spatial origins (Fig 2b). In our simulation data, four types of cells can be classified as rare: 2-Mesothelium and Submucosal Secretory have only 1 cell each, Myofibroblasts has 2, and Fibromyocytes has 7. The boxplots indicate that our approach successfully assigned all these rare cell types to their correct spatial positions.

**Figure 2:**
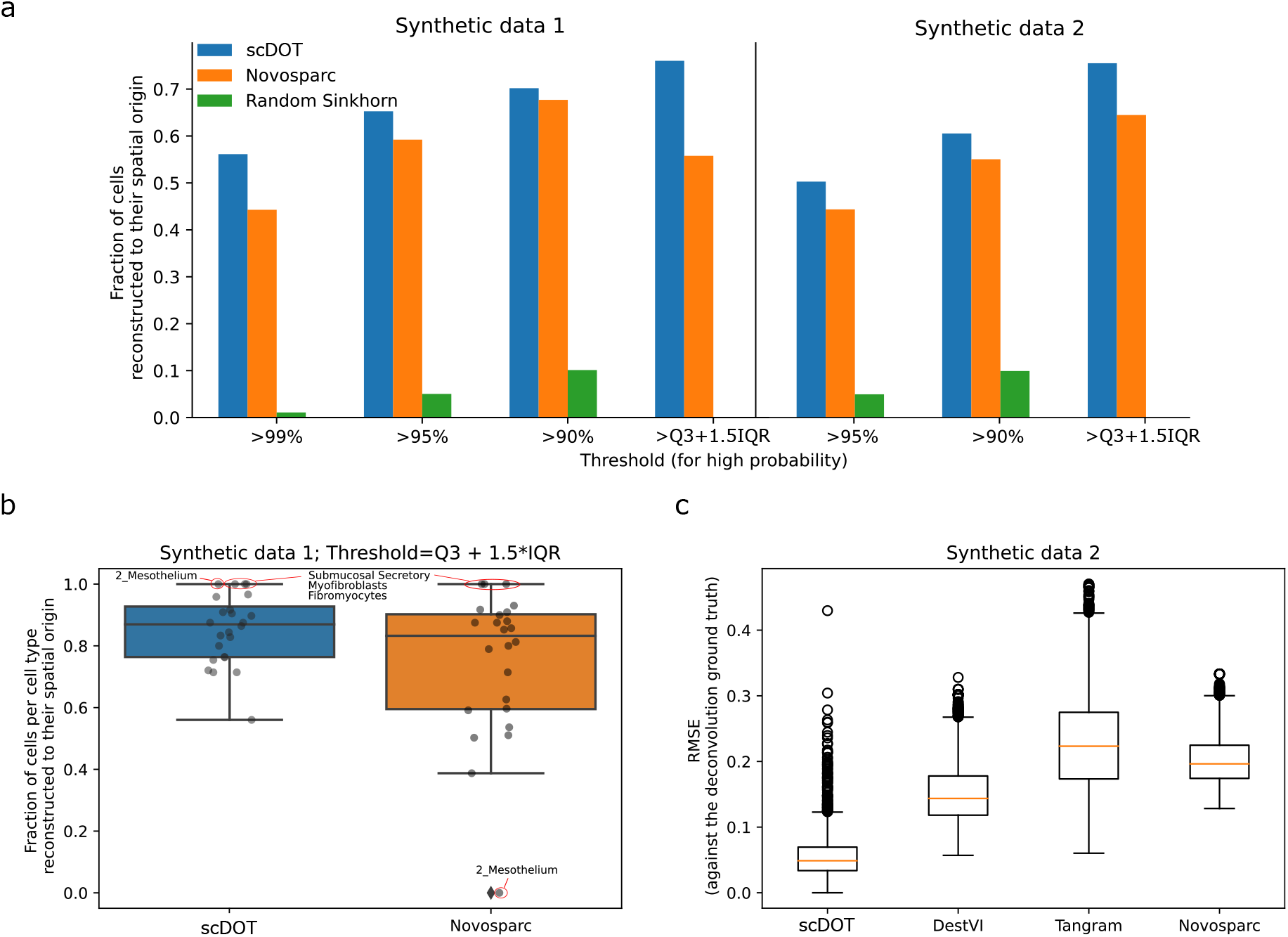
Performance on synthetic datasets. (a) OT results of simulation datasets 1 and 2 demonstrate that by using different thresholds to define a high probability, we can assign nearly 80% of cells to their spatial origin. scDOT was benchmarked against two other methods: Novosparc, a spatial reconstruction method based on Gromov-Wasserstein distance, and Random Sinkhorn, a naive method that learns the optimal transport coupling with a random cost matrix. The results demonstrate the superior performance of scDOT in all cases. (b) Detailed results of simulation data 1 (with a threshold higher than the 3rd quantile plus 1.5 times the IQR) highlight the effectiveness of scDOT and spatial mapping methods in general for rare cell types. The boxplots illustrate the fraction of correctly reconstructed cells per cell type. Each point represents a single cell type (*c* = 24). Among the considered rare cell types (2-Mesothelium and Submucosal Secretory with 1 cell, Myofibroblasts with 2 cells, and Fibromyocytes with 7 cells), scDOT successfully mapped these rare cell types to their exact spatial locations (fraction = 1.0), while Novosparc failed to map 2-Mesothelium to its spatial location (fraction = 0.0). (c) The root-mean-square-error (RMSE) of the deconvolved cell-type proportions compared to the ground truth is evaluated for synthetic data 2, consisting of 9 cell types across 3072 spots. scDOT, along with other methods including DestVI, Tangram, and Novosparc, is compared in terms of RMSE. The boxplots demonstrate that scDOT outperforms the other methods, as indicated by the lower RMSE values. The boxplots display the median (middle line), 25th and 75th percentiles (box), and 5th and 95th percentiles (whiskers).

### 2.2 Comparison to other methods on spatial mapping and cell type deconvolution

#### Spatial mapping

We evaluate the performance of scDOT in spatial mapping and compare it with other existing methods. Figure 2a presents the results for Synthetic data 1, where the threshold is set above Q3 + 1.5×IQR. scDOT achieves the highest outcome at this threshold, while the outcome of Novosparc is drastically decreased compared to the outcome at thresholds above the 90th and 95th quantiles. This observation suggests that our probabilistic mapping exhibits a heavier-tailed characteristic, which is a more desirable property for accurate spatial mapping.

Furthermore, we find that the reconstruction results are influenced by the dataset used. For Synthetic data 2, scDOT achieves a high outcome when the threshold is set above Q3 + 1.5×IQR, with 76% of cells successfully reconstructed. However, stricter thresholds lead to a more rapid decay in the outcomes, with only 50% of cells being reconstructed at the threshold above the 95th quantile. Nevertheless, across all cases, scDOT consistently outperforms both Novosparc and the naive baseline of Random Sinkhorn.

In terms of accurately mapping rare cell types to their spatial positions, scDOT successfully assigns all four rare cell types with a fraction of 1.0. However, Novosparc failed to accurately map 2-Mesothelium to its spatial location, as indicated by a fraction of 0.0. Also, as indicated in Figure 2a, scDOT mapped 76% cells correctly while Novosparc mapped 56% cells correctly; these 20% differences is not shown in Figure 2b since the difference in the number of cells per cell type is not considered.

#### Deconvolution

To benchmark the results of cell type deconvolution, we applied scDOT to synthetic data 2 and compared it with three other methods: DestVI [29], Tangram [3], and Novosparc [37]. The synthetic dataset comprised nine cell types distributed across 3072 spots. We specifically chose these three deconvolution methods as they represent distinct computational techniques tailored for spatial transcriptomics data. DestVI is a probabilistic-based method, Tangram utilizes deep learning, and Novosparc is an OT-based method. All three methods require spatial transcriptomics data as input and scRNA-seq data as a reference. Comparing the root-mean-square-error (RMSE) of the deconvolved cell type proportions with the ground truth, scDOT outperformed the other three methods (see Figure 2c). The mean RMSE scores for scDOT, DestVI, Tangram, and Novosparc were 0.06, 0.15, 0.23, and 0.20, respectively. It’s worth noting that Novosparc is not designed for direct computation of cell type deconvolution but rather for mapping cells to spots. As a result, the deconvolution results are calculated by multiplying the coupling matrix *γ* with the cell-by-cell type relation matrix *C*, i.e., *P* = *γ* × *C*.

### 2.3 Identfiying the spatial patterns of the distribution of specific cell types

We used paired IPF scRNA-Seq and spatial dataset to test the ability of our mapping method to infer cell-cell interactions (Figure 3). Among the 29 cell types (Methods), Multiciliated, Secretory and Basal cells exhibited prominent and distinct spatial patterns. Notably, Multiciliated, Secretory, and Basal cells were found to be in close proximity to each other, both in the upper lobe and lower lobe of the tissue. This observation aligns with the traditional view of the airway epithelial mucosal layer, which incorporates basal cells in close proximity to secretory and ciliated cells, forming a tight unit. This unit serves as a physical barrier while remaining responsive to the inhaled environment through interactions with submucosal fibroblasts, smooth muscle cells and cells and molecules from the immune system [18].

**Figure 3:**
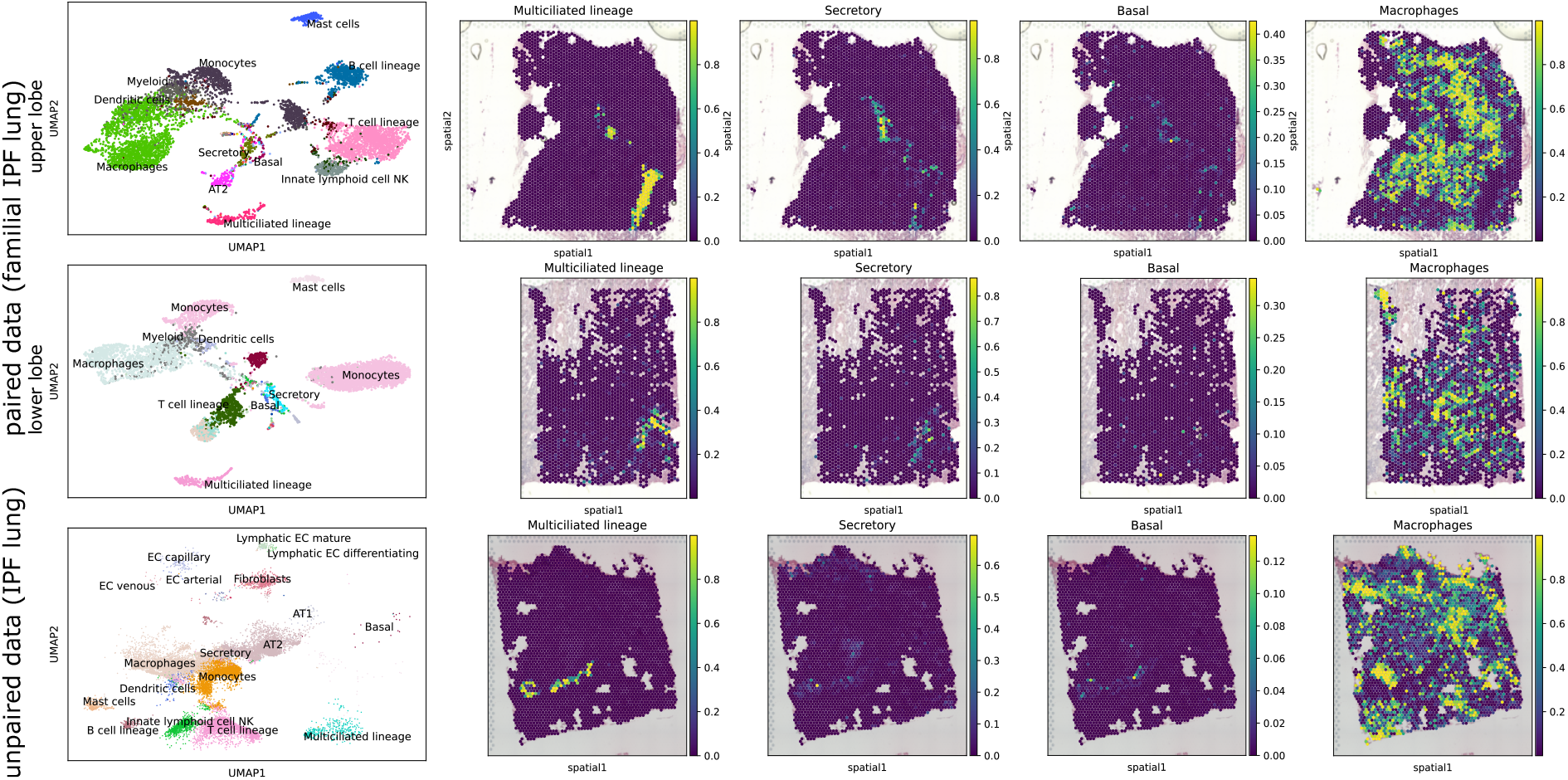
Spatial distribution patterns of multiciliated, secretory, basal, and macrophage cells across different datasets. **Top**: A UMAP representation of scRNA-seq data, along with the spatial patterns of the selected cell types in the upper lobe slice of the paired familial IPF lung. **Middle** A UMAP representation of scRNA-seq data and the corresponding cell types in the lower lobe slice of the same sample. **Bottom** A UMAP representation and spatial distribution of selected cell types in the unpaired IPF lung sample. Notably, multiciliated, secretory, and basal cell types exhibit distinct and prominent spatial patterns. Importantly, these cell types consistently exhibit close proximity to each other across all three datasets, consistent with previous studies on the organization of the respiratory system [18, 9, 27].

Secretory and multiciliated cells are known to be located in close proximity to each other within the respiratory tract, including the lungs. Together, they form a self-clearing mechanism that efficiently removes inhaled particles from the upper airways, preventing their transfer to deeper lung zones [9]. The coordinated action of multiciliated cells, with their motile cilia, and secretory cells, responsible for mucus production and secretion, enables the effective clearance of inhaled particles and maintains the integrity of the respiratory system [27].

Basal cells, positioned closer to the basement membrane, further contribute to the organization and functioning of the airway epithelium. They provide structural support and are responsible for the regeneration and repair of the airway epithelial layer [18].

The spatial organization of Multiciliated, Secretory, and Basal cells in close proximity to each other emphasizes their interdependence and coordinated functioning in maintaining the respiratory barrier and facilitating efficient clearance mechanisms. This finding underscores the significance of the spatial arrangement and interactions of diverse cell types within the airway epithelium for the overall homeostasis and defense of the respiratory system.

Conversely, immune cell types such as Macrophages and T cells lineage, which were characterized by a larger number of cells, displayed a more scattered distribution throughout the tissue. Yet, the spatial distribution of these two cell types are complementary to some degree (Figure 3 and 4, Supplementary figures), reflecting the fact that they are both important components of the immune system and play complementary roles in defending against infections and maintaining immune homeostasis. On the other hand, cell types with a smaller cell count, such as smooth muscle (consisting of only 2 cells in total), exhibited a spatial arrangement in adjacent spots (Supplementary figures).

**Figure 4:**
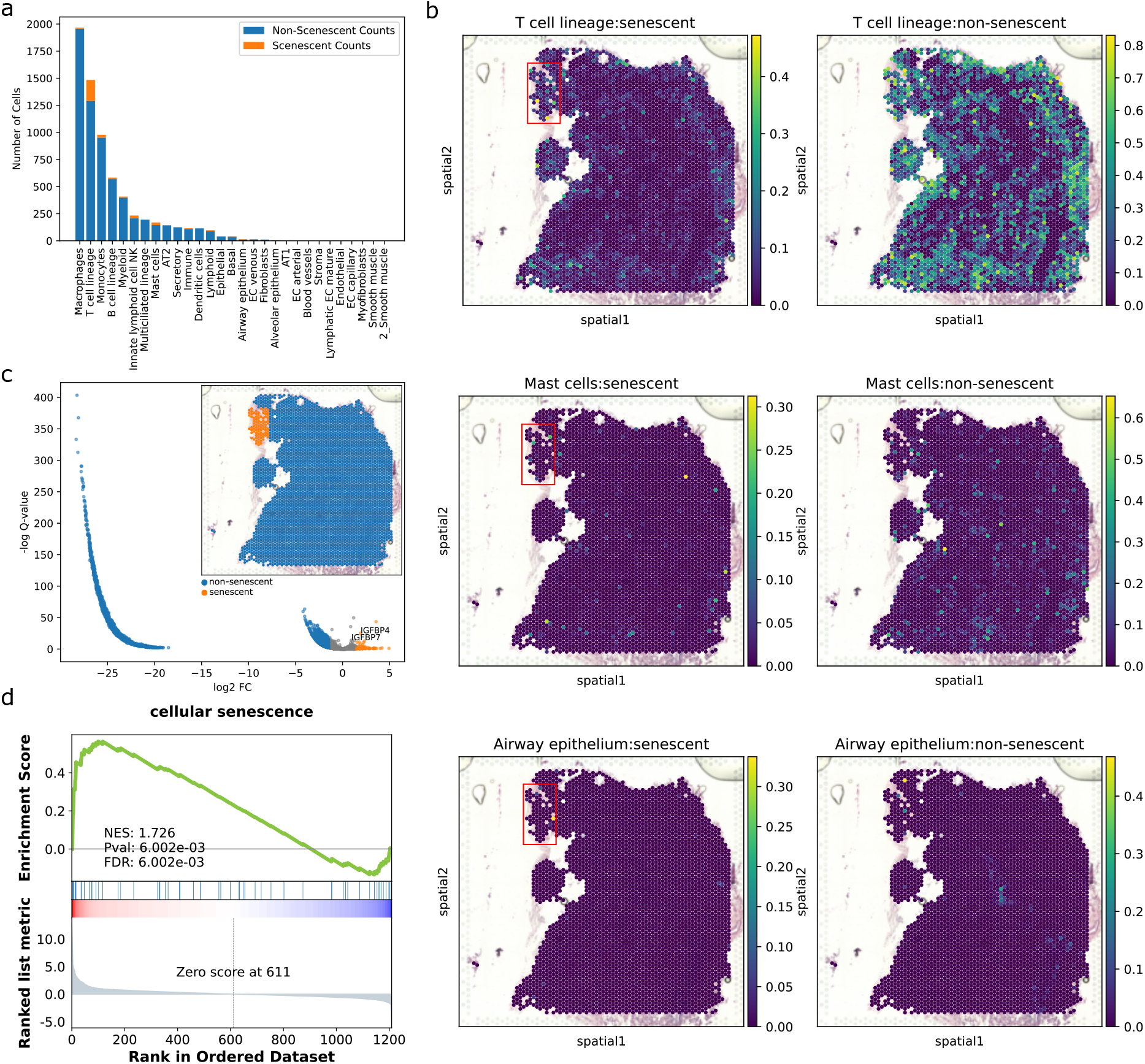
Analysis of cellular senescence reveals the spatial collocation of senescent cells. (a) The number of senescent cells and non-senescent cells for each cell type is depicted. T cell lineage, mast cells, and airway epithelium exhibit the highest fraction of senescent cells. (b) Spatial distribution of senescent and non-senescent cells for the three aforementioned cell types. Notably, the three different senescent cell types are spatially collocated in the upper left corner of the tissue. (c) Differentially expressed genes for the manually annotated senescent region (colored in orange) in the upper left corner of the tissue (as depicted in panel (b) of this figure and the upper right corner of this panel). Among the top-ranked DEGs are IGFBP4 and IGFBP7, which are also senescent marker genes. (d) Gene set enrichment analysis (GSEA) plot. The top-ranked DEGs (as shown in panel (c) of this figure) are enriched in the gene set consisting of 340 senescent marker genes.

These patterns were also observed in the unpaired data, particularly with regards to the multiciliated lineage and secretory cell types (Figure 3), demonstrating the generality of our approach on unpaired datasets.

### 2.4 Cell-cell proximity analysis

To quantitatively illustrate the spatial distribution and proximity of multiciliated, secretory, and basal cells described in section 3.3 of this paper, we employed the neighborhood enrichment score. This score between two cell types represents the z-score derived from a permutation test that tallies the neighboring spots consisting of either cell type. Consistent with the spatial patterns depicted in section 3.3 and Figure 3, we observed the highest enrichment score between the multiciliated lineage and itself across various datasets (69.46 in the upper lobe of familial IPF paired data, 29.31 in the lower lobe of the same data, and 47.98 in the IPF unpaired data). The score between Multiciliated and Secretory cell types is also one of the highest (19.40 in the upper lobe of the paired dataset, 12.25 in the lower lobe, and 5.06 in the unpaired dataset). In contrast, the scores between Macrophages and T cells are among the lowest across datasets, with scores of −25, −5.75, and −15.83 in the upper lobe, lower lobe, and unpaired dataset, respectively. These scores reflect the fact that they are complementary, as indicated in section 3.3 (see Supplementary Figure 1). It is important to note that the neighborhood enrichment scores were estimated at the spot-level and only considered the dominant cell type of each spot, which is defined as the cell type with the highest proportion within that particular spot.

At the cell level, we constructed a cell-cell spatial proximity graph based on OT placement (see Methods). The graph was then summarized by cell types, quantifying the physical proximity between each cell type by counting the direct neighboring cells within the same type (see Supplementary Figure 1d and e, Supplementary Table 1). Once again, the multiciliated lineage exhibited the highest normalized counts with itself across datasets, consistent with the results obtained from the enrichment score and described in Section 3.3. In the paired dataset, basal and secretory cells also demonstrated a strong association with the airway epithelium, providing additional evidence for the spatial organization of the respiratory system as discussed in Section 3.3. In contrast, immune cells such as T cells and macrophages displayed connections to various cell types, reflecting their dispersed distribution throughout the tissue. Notably, in the IPF lung sample, fibroblast cells exhibited a distinct spatial pattern and were found to be in close proximity to 2-smooth muscle cells and myofibroblast cells, supporting previous research suggesting that *α* smooth muscle actin-expressing fibroblasts, referred to as myofibroblasts, serve as markers of progressive lung injury and play a central role in detrimental remodeling and disease progression [41, 20] (Supplementary Figure 1, Supplementary Table 1, Section 3.6).

### 2.5 Identification of senescent markers

For cellular senescence analysis, we profiled two new spatial datasets. The first included paired scRNA-Seq data from a familial IPF lung sample, and the other consists of unpaired data from an IPF lung sample (Methods).

#### Paired data of familial IPF lung sample

We first identified in the scRNA-seq data, cell types with a large fraction of cells exhibiting senescent. For this, we used a list of 68 senescent marker genes (*Methods*). Within each cell type, we separated the cells into senescent and non-senescent cells (Figure 4a, b). For this familial IPF lung sample, the ratio of senescent cells to non-senescent cells is low. For most cell types we observed very few senescent cells. For other we found more. For example, for Mast cells, T cell lineage, and Airway epithelium we identified 14%, 13%, and 17%, respectively. We thus focused on these three cell types. for these we had 24, 193, and 3 senescent cells for Mast cells, T cell lineage, and Airway epithelium, respectively. Next, we manually annotated the regions where senescent cells from different cell types are collocated (Figure 4b, c). For these regions we computed differentially expressed genes (DEG) w.r.t. the rest of the tissue. As expected, given the way we selected these regions we found among the top ranked DEG IGFBP4 and IGFBP7 (t-test p-values are 1.1e-11 and 7.2e-07 respectively), which are both senescent marker genes (Figure 4d). We next performed gene set enrichment analysis (GSEA) with this ranked gene list and a gene set of 340 senescent markers (which is a superset of the 68 senescent marker genes set we used for re-annotation, Supplementary Data 1), we confirmed that cellular senescence is enriched–with p-value = 0.006002; FDR = 0.006002, and the normalized enrichment score is 1.726–in the annotated region (Figure 4d). The leading-edge subset of genes in this analysis comprised IGFBP4, IGFBP7, FGF7, THBS1, IGF1, IGFBP6, IL6, SERPINE2, PIM1, ALDH1A3, SERPINE1, COL1A2, ANGPTL4, CYP1B1, and PLAU.

While IGFBP4 and IGFBP7 belong to the initial set of 68 senescent marker genes, the remaining genes are part of the larger set of 340 senescent marker genes. Of particular note, IGFBP4 and IGFBP7 are SASP factors that have been identified as key components needed for triggering senescence in young mesenchymal stem cells (MSC) [42]. The pro-senescent effects of IGFBP4 and IGFBP7 are reversed by single or simultaneous immunodepletion of either proteins from the conditioned medium (CM) from senescent cells [42]. According to a previous study, prolonged IGF1 treatment leads to the establishment of a premature senescence phenotype characterized by a unique senescence network signature [34]. Combined IGF1/TXNIP-induced premature senescence can be associated with a typical secretory inflammatory phenotype that is mediated by STAT3/IL-1A signaling [34].

### 2.6 Inferring Cell-Cell interactions driving senescence

We also looked at the cell type neighborhood of senescent cells. These are summarized in Figure 5a. We observe that senescent cells are often close to non-senescent cells of the same type (e.g., senescent T cells to non-senescent T cells) which can explain why some cell types have a much higher percentage of senescent cells than others.

**Figure 5:**
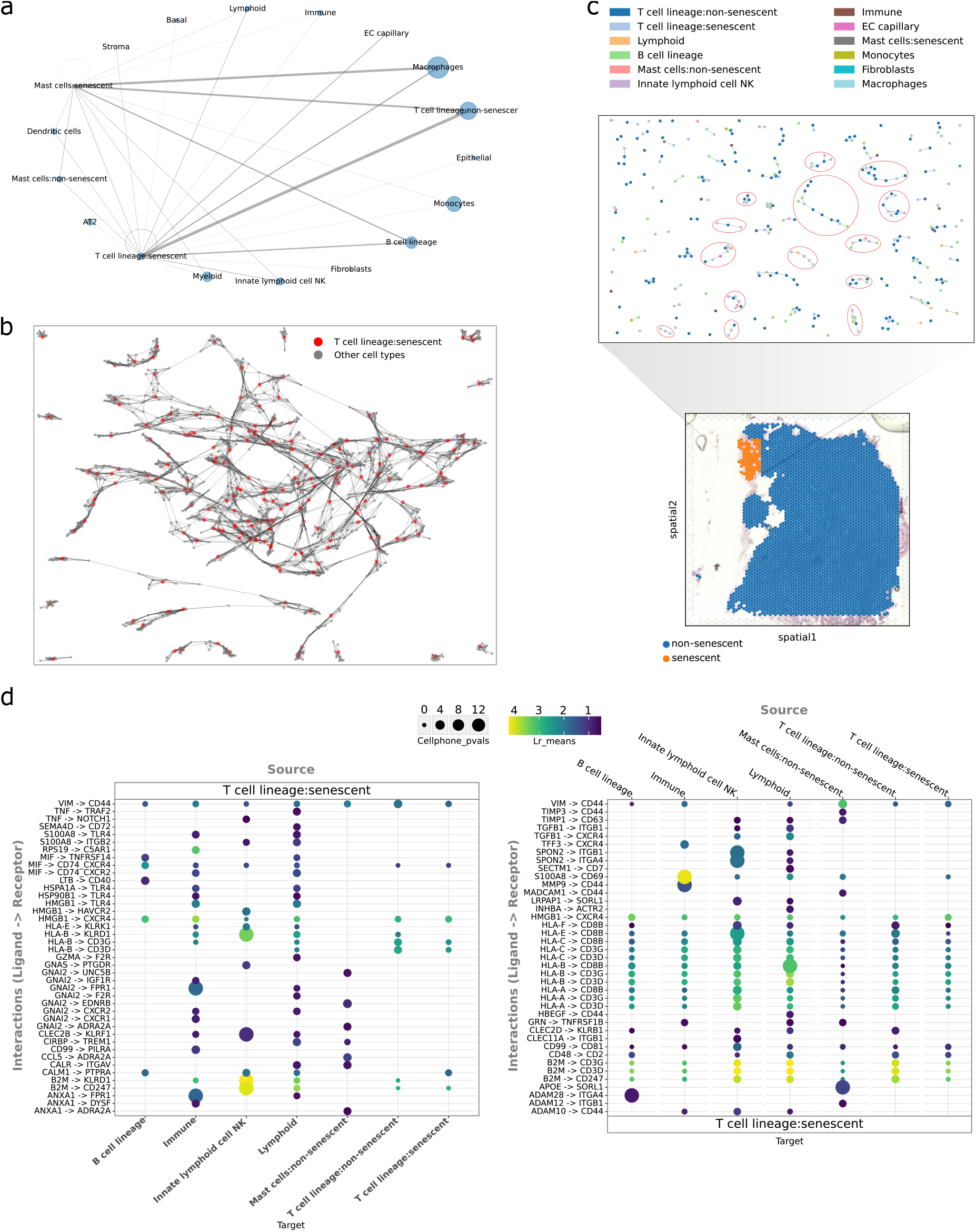
Analysis of senescent cell-cell communication in the upper lobe of the familial IPF lung sample. (a) The graph summarizes the spatial neighborhood of senescent mast cells and T cell lineage. Nodes represent cell types, and edges indicate direct neighboring relations in physical proximity. The size of each node corresponds to the number of cells within a cell type, while the width of the edges represent the number of neighboring cells of a specific cell type (i.e., the total node degree per neighboring cell type). Edges representing a small number of neighbors are omitted. As can be seen, senescent cells are close to both non-senescent cells within the same cell types and senescent cells belonging to different cell types. (b) Cell-cell spatial neighborhood of senescent cells for the T cell lineage. The validity of this neighborhood graph is assessed in Supplementary Analysis. (c) The subgraph of the cell-cell neighborhood depicted in panel (b), specifically showing the cells located in the senescent region (colored orange). (d) The results from CellphoneDB display the co-expressed ligand-receptor pairs between senescent cells of the T cell lineage and all other cells within the subgraph illustrated in panel (c).

Utilizing the CellPhoneDB [11], we further identified the ligand-receptor (LR) pairs involved in the cell-cell interactions within the neighborhood of senescent cells (i.e., within the graph *G*^′^) (Figure 5d). We observed that 11 senescent markers, namely B2M, CALR, CCL5, CD44, HMGB1, IGF1R, MIF, TNF, VIM, MMP9, and TNFRSF1B, were significantly overrepresented in the list of ligands and receptors identified by CellPhoneDB (hypergeometric test p-value = 0.00072). Among the LR pairs involved in senescent-to-senescent cell-cell communication (i.e., between senescent T cells), most of the pairs include senescent marker genes. The other remaining LR pairs involve the HLA gene family (which is essential for T cell activation). For example, HLA-E acts as an inhibitory signal for NK and CD8 T cells—and depletion of HLA-E renders senescent cells susceptible to elimination by both NK and CD8 T cells [39]. Another LR pair involves S100A8, which increases with age, inducing inflammation and cellular senescence-like phenotypes in oviduct epithelial cells [35, 14].

#### Unpaired data from IPF lung sample

To demonstrate the general utility of the method for unpaired data, we performed the same analysis as described for the paired data mentioned above for another spatial dataset we profiled, this time without matched scRNA-Seq (Methods). Using a scRNA-seq dataset of an IPF lung sample, we were still able to identify several of the same senescence cell types as in the paired dataset, including T cells and mast cells. There were 300 assigned senescent cells out of the total 3747 T cells and 11 assigned senescent mast cells out of the total 249 mast cells. We also observed high fraction of senescence cells for other cell types including for fibroblasts (290 out of the total 461 fibroblast cells) and 2-smooth muscle (8 out of 21).

We again observed that senescence cells co-localized in the same regions (Figure 6a). While T cells tended to be distributed throughout the tissue, there is a high fraction of senescent cells co-localized with fibroblasts and mast cells (Figure 6a). Fibroblasts and 2-smooth muscle cells co-localized in specific regions, with a total of four overlapping regions as depicted in Figure 6a. Since senescent cells tend to co-localize with other cells of the same type, most senescent fibroblast cells and 2-smooth muscle cells also co-localized (except for the region in the upper left corner of the tissue, which exhibited only senescent fibroblast cells). These observations of senescent spatial distribution align with previous studies suggesting that senescent cells have the potential to influence neighboring cells through processes collectively referred to as the senescence-associated secretory phenotype [31].

**Figure 6:**
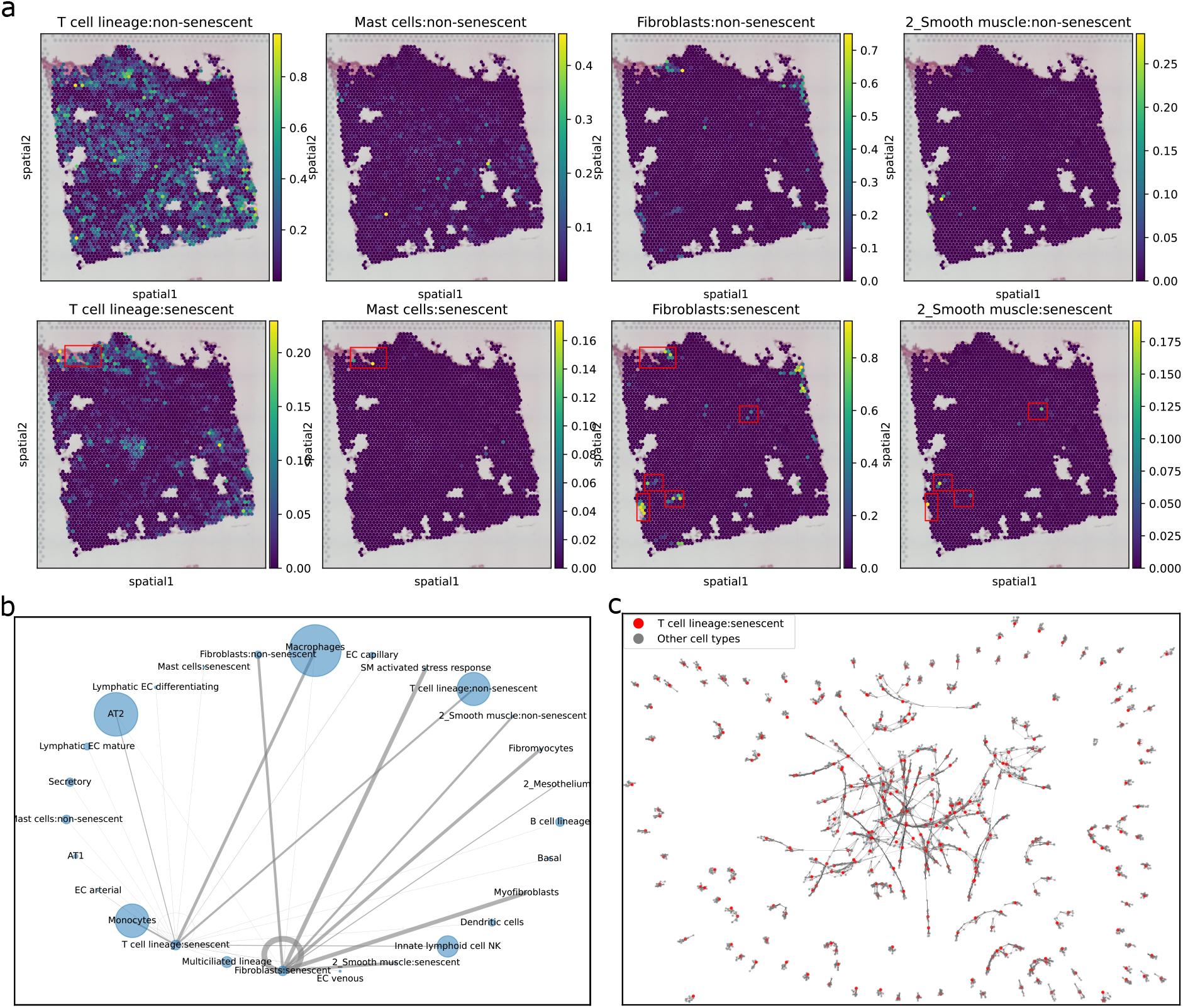
The analysis of senescent cell-cell neighborhood for unpaired IPF dataset. (a) Spatial distribution of senescent and non-senescent cells for T cells, Mast cells, Fibroblasts, and 2-Smooth muscle. Red rectangles indicate regions where senescent cells of multiple types are co-located. (b) The graph summarizes the spatial neighborhood of senescent Fibroblasts and T cells. Nodes represent cell types, and edges represent direct neighboring relations in physical proximity. The size of the nodes corresponds to the number of cells within a cell type, and the width of the edges corresponds to the number of neighboring cells of a specific cell type (i.e., the total node degree per neighboring cell type). Edges representing a small number of neighbors are omitted. The graph demonstrates that senescent cells are neighbors to non-senescent cells within the same cell types, as well as to senescent cells belonging to different cell types. (c) The cell-cell spatial neighborhood graph of senescent cells of the T cell lineage. The validity of this neighborhood graph is assessed in Supplementary Analysis.

Figure 6b and c illustrate the physical proximity among cells of different cell types. Similar to the paired data of the familial IPF lung sample, senescent cells are closely clustered together and near cells of the same type. As shown in Figure 6b, the senescent T cells are adjacent to other T cells, mast cells, and macrophages. The cell-to-cell spatial neighborhood graph, with nodes representing senescent T cells and their immediate neighbors, is depicted in Figure 6c. The validity of this neighborhood graph is assessed in Supplementary Analysis. For a more specific focus on senescent fibroblasts, a cell-to-cell neighborhood graph can be found in Supplementary Figure 2.

## 3 Discussion

In this study, we introduced a novel method for integrating single-cell and spatial transcriptomics, addressing the simultaneous tasks of cell type deconvolution and spatial reconstruction. The challenge of spatial reconstruction lies in the non-linear relationship between gene expression profiles of single-cells and the spatial transcriptomics data [46], as well as the inherent uncertainty in high-resolution mapping. However, by incorporating internal references from cell type deconvolution, we can modulate and enhance the precision of this task.

Our method, scDOT was shown to efficiently and accurately assign individual cells to their spatial origins using synthetic data. By combining OT and deconvolution scDOT improves on all prior methods we compared to. We also used scDOT to study and analyze new paired and unpaired spatial transcriptomics data from IPF and familial IPF lungs. We observed that senescent cells tend to co-localize in specific regions and are in close proximity to cells of the same type. While the distribution of senescent T cells appears sparse in both datasets, we noted a denser population of senescent fibroblast cells in the IPF lung compared to the familial IPF lung, which can be explained by the paracrine senescence and is consistent with previous studies indicating that senescent fibroblasts contribute to the pathogenesis of IPF through various mechanisms [1, 48, 26].

The integration of single-cell and spatial transcriptomics has been a topic of interest in recent years [28], with a number of multiview learning approaches suggested [36]. A crucial aspect of this integration is assessing the similarity of gene expression levels between cells and spatial spots. Unlike prior methods that utilized optimal transport, which rely on fixed cost matrices to represent the dissimilarity between cells and spots, scDOT utilizes a differentiable optimization layer in a deep declarative network to dynamically learn the cost matrix [16]. This use of optimal transport can be formulated as a domain adaptation problem, and the learned cost matrix holds potential for further applications involving mass transportation between the two modalities of other types of data.

Comparative studies and benchmarks exist for cell type deconvolution in spatial transcriptomics data [23, 24, 50]. Since there is no universal evaluation metric that applies to all scenarios, comparisons among methods depend on datasets and evaluation metrics used, such as root mean square error and Lin’s concordance correlation coefficient, which may not consistently correlate [23, 6]. In our paper, we compared our method with recent approaches representing computational techniques like deep learning, probabilistic modeling, and optimal transport. While the performance of these methods may vary, certain high-performance methods, particularly Tangram [3], have been reported [23, 24, 50]. Additionally, note the normalization of our synthetic data 2, making methods utilizing count matrices as input, such as Stereoscope [2] and Cell2Location [21], inapplicable.

An important component of our biological analysis focused on IPF and familial IPF lung tissue was the identification of senescent cells. Evaluating cellular senescence poses challenges as there are various approaches, such as assessing senescent gene markers or morphological features of senescent cells. Additionally, different cell types or diseases may require distinct sets of senescent markers due to the complex nature of the senescence process. In our study, we employed a combined list of senescent markers and categorized cells within each cell type as either senescent or non-senescent. However, senescent states can exist on a continuum, ranging from non-senescence to primary senescence, and different markers may be associated with primary and secondary senescence. Still, using scDOT we were able to identify cell-cell spatial neighborhood, which can aid in assessing senescent cells in close physical proximity. It also allowed us to explore how senescent cells reorganize and impact their environment and nearby cells. Cells neighboring senescent cells can transition into a secondary senescent state. Hence, the influence of senescence can be approached as a diffusion problem within a network, where cells reach a senescent state through contact with senescent neighbors. This network-based diffusion approach, relying on the spatial mapping of individual cells to their origins, holds promise for fruitful future investigations. scDOT is implemented in PyTorch and is available for download from https://github.com/namtk/scDOT.

## 4 Methods

### 4.1 Data sets

To investigate the effects of the proposed method that combines cell-type deconvolution and spatial reconstruction, we collected both synthetic and real data. Since there is no immediate method to assess the performance of cell-type deconvolution and spatial reconstruction tasks on real data, we generated two simulation datasets to evaluate and benchmark scDOT as well as other related methods against the ground truth. It is important to note that, for benchmarking the deconvolution task, methods designed for spatial reconstruction can be utilized. However, for benchmarking the reconstruction task, methods solely designed for cell-type deconvolution cannot be used, as inferring the fine-grained mapping *γ* of individual cells from a coarse-grained mapping *P* of cell clusters poses a challenging inverse problem, even though inferring the cell type proportion *P* from the coupling matrix *γ* is straightforward (*P* = *γ* × *C*).

#### 4.1.1 Synthetic data sets

##### Synthetic data set 1

The synthetic data 1 is generated based on Gaussian Process (GP) by assuming that the nearby spots have similar proportions of cell types as well as gene expressions [29]. Here, we used scRNA-seq data of an IPF lung tissue and projected the cells from this data onto grids, which represent the spatial coordinates obtained from a different IPF lung sample’s upper lobe lung slice. Thus, scRNA-seq data is real while spatial locations are synthetic for this dataset. See Supporting methods for more details.

##### Synthetic data set 2

For the synthetic data 2, we conducted simulations using gene expression data from individual cells obtained through multiplex error-robust fluorescence in situ hybridization (MERFISH) in the mouse medial preoptic area (MPOA) [32, 33]. By aggregating the gene expression information of cells within spatially contiguous pixels, we created a representation of the spatial organization. See Supporting Methods for more details.

#### 4.1.2 Real data sets

##### Preparation and data collection of single-cell RNA sequencing and spatial transcriptomics

Tissue samples were obtained by the Human Tissue Biorepository at The Ohio State University from the explanted lungs of patients diagnosed with idiopathic pulmonary fibrosis (IPF) and familial IPF after a Total Transplant Care Protocol informed consent and research authorization from the patient. The tissue biorepository operates in accordance with NCI and ISBER Best Practices for Repositories.

###### For single-cell RNA sequencing (scRNA-seq)

Samples of 15 g of upper and lower lobe lung parenchyma tissue were washed with PBS, minced finely with scalpels, and digested using an enzyme cocktail (1 mg/mL of liberase DL, DNase I, DMEM) for 2 hours at 37°C with rocking. Cell suspension was filtered through a serial filter of 300 *μ*m, 100 *μ*m, and 70 *μ*m strainers. After straining, the cell suspension was centrifuged at 500g for 7 minutes, the supernatant was removed, and 1x RBC lysis buffer was added to the pellet and incubated at 4°C for 7 minutes and then filtered through a 40 *μ*m strainer to remove the agglomerated dead cells. Finally, cell number and viability were determined using a countess automatic cell counter (Invitrogen). Whole lung cell suspension was loaded on the Chromium Controller, according to 10x Genomics protocol. 3’ Gene Expression libraries were sequenced on Illumina sequencer with read lengths of 28 cycles Read 1, 10 cycles i7 index, 10 cycles i5 index, 90 cycles Read 2. ScRNA-seq data was extracted from the raw sequencing data using Cell Ranger (version 7.1.0, 10x Genomics).

###### For spatial transcriptomics

Tissue sections of ≤ 6.5 x 6.5 mm from the upper and lower lobe of lung parenchyma were used for spatial analysis. After collection, samples were fixed for 24 hours in 10% neutral buffered formalin and embedded in paraffin (wax) to create a formalin-fixed paraffin-embedded (FFPE) block. Sections of 5*μ*m were then cut from the FFPE blocks onto Visium slides (10x Genomics) and processed according to the manufacturer’s protocol. Scan of H&E staining was performed with EVOSTM M7000 microscope (Invitrogen) using a 10x objective. FFPE libraries were prepared according to 10x Genomics protocol and sequenced on Illumina sequencer to a read depth of at least 25k reads/spot, with read lengths of 28 cycles Read 1, 10 cycles i7 index, 10 cycles i5 index, 50 cycles Read 2. Spatial transcriptomics data was extracted from the raw sequencing data using Space Ranger (version 2.0.0, 10x Genomics).

##### Paired familial IPF lung data set

We obtained two paired datasets of single-cell and spatial transcriptomics from a patient with familial IPF lung, one of which is from the upper lobe slice and the other from the lower lobe slice. The upper lobe pair contains 6762 cells and 3336 spots while the lower lobe pair contains 6173 cells and 2246 spots. For each of these two paired datasets, we preprocessed the data by (1) removing lowly expressed genes of both two data modalities, keeping genes that have at least 10 counts, and (2) removing cells with low counts, keeping cells that have at least 500 counts and 500 genes expressed, then (3) obtaining the common gene sets for both modalities by taking the intersection of the two gene sets.

The cell type annotations were transferred from the Lung cell atlas (HLCA) using scArches and FastGenomics platform.

To re-annotate cells that reflect senescent states, we utilized a list of 68 senescent markers (Supplementary Data 1), then calculated the average expression of the marker genes across all cells. Next, senescent cells were identified as having a higher than 95 percentile of average expression of the marker genes.

##### Unpaired IPF lung data set

To demonstrate the general utility of the method even for non-paired data, we obtained an unpaired scRNA-seq and spatial transcriptomics dataset from two different IPF patients. While the preparation for spatial transcriptomics is the same as for the paired data, the preparation for single-cell RNA sequencing is described as follows. Single-cell sequencing of human lung tissue was performed as previously described [44, 17]. In short, human lung tissue (IPF) was homogenized, and 4 g of tissue were digested by dispase/collagenase (Collagenase: 0.1U/mL, Dispase: 0.8U/mL, Roche) for 1 hour at 37°C. Samples were successively filtered through nylon filters (100 *μ*m and 20 *μ*m) followed by a percoll gradient. Single epithelial cell suspensions were loaded onto a Chromium single-cell chip (Chromium™Single Cell 3’ Reagent Kit, v2 Chemistry) to obtain single-cell 3’ libraries for sequencing. cDNA obtained after droplet reverse transcription was amplified for 14 cycles and analyzed using Agilent Bioanalyzer. The barcoded libraries were sequenced using Illumina NextSeq-500 through the University of Pittsburgh Genomics Core Sequencing Facility, aiming for 100,000 reads per cell and capturing 10,000 per library.

The single-cell data contains 25,260 cells, and the spatial data, which consists of an upper lobe lung slide, contains 3,412 spots. The preprocessing, cell type annotation, and senescence re-annotation were carried out following the same procedures as for the paired familial IPF lung dataset.

It is important to note that, since the familial IPF lung datasets are paired, the coordinates of cells after spatial reconstruction represent the actual tissue coordinates. However, for the unpaired IPF lung dataset, the inferred cell coordinates do not directly reflect the actual tissue coordinates. Instead, they serve as an intermediate step to infer the relative spatial relationships among cells.

### 4.2 Cell type deconvolution

For gene expression, cell type deconvolution can be formulated as a nonnegative least squares (NNLS) problem, where the goal is to estimate the relative abundances of different cell types by solving for the nonnegative coefficients of a linear combination of their respective gene expression profiles. Specifically, a multicellular resolution spatial transcriptomics profile *Y* ∈ ℝ^*m*×*p*^ of *p* genes across *m* spots each of which contains transcripts from multiple cells can be represented as *Y* = *PS* in which *P* ∈ ℝ^*m*×*c*^ is the cell type proportions to be estimated and *S* ∈ ℝ^*c*×*p*^ is the signature matrix consisting of known gene expression profiles for each cell type of the total *c* cell types. We solved for *P* the following nonnegative least squares problem:

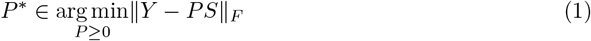

There are several solvers available for solving a NNLS problem, including Lawson-Hanson’s active set method [22]. Here, we used projected gradient descent [25].

### 4.3 Mapping single cell to spatial images

The spatial reconstruction task involves assigning cells from scRNA-seq data to a predicted corresponding location in a tissue sample. Note that such assignment, implicitly, also provides deconvolution of the spot data assuming that the cell types for cells in the scRNA-Seq data are known. Here we formulate this as an optimal transport (OT) problem from scRNA-seq dataset *X* ∈ ℝ^*n*×*p*^ of *p* genes across *n* cells to spatial transcriptomics dataset *Y* ∈ ℝ^*m*×*p*^ of *p* genes across *m*. OT is commonly used to model the coupling between two probability distribution. In our case we use it to model the transport of gene expression from one dataset to another in an optimal way. By solving the optimal transport problem, it is possible to estimate the optimal coupling and quantify the degree of similarity between the datasets. Formulating the spatial reconstruction task as an optimal transport problem involves constructing a cost matrix 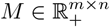 representing distances between cells of *X* and spots of *Y*. Here, we used cosine distance 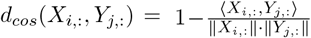, which is scale-invariant and can account for differences in measurement sensitivity between the two technologies. Furthermore, scale-invariant cosine dissimilarity is well-suited for handling the fact that expression of a spot in the spatial transcriptomics dataset is the mixture or sum of multiple cells in the scRNA-seq dataset. Specifically, the coupling matrix 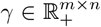 is solved for obtaining the optimal transport as follows:

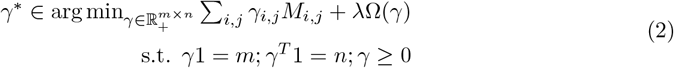

where 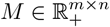 is the cost matrix defining the cost of moving gene expression from cell *a*_*i*_ to spot *b*_*j*_ and Ω(*γ*) =∑_*i,j*_ *γ*_*i,j*_ log (*γ*_*i,j*_) is an entropic regularization term [8]. The entropic regularization version of optimal transport can be solved by Sinkhorn-Knopp’s alternative projection algorithm [8]. In other words, this minimization process aims to match cells with similar expression profiles to spots with similar transcriptomic characteristics, measured by cosine similarity, thereby capturing the underlying biological relationships between the two datasets.

It is important to note that we utilized the entropy regularization version of optimal transport, resulting in a probabilistic mapping between cells and spots. This probabilistic coupling, represented by the left-stochastic matrix *γ*, indicates the likelihood of a specific cell being associated with a particular spot. This probabilistic coupling offers computational efficiency and eliminates assumptions about the number of cells in a spot, including cases where a cell may reside on the boundary of two spots.

### 4.4 Combination of deconvolution and mapping

OT for spatial and scRNA-Seq data is challenging since spatial data is often sparse leading to less dependable inferred individual cell-spot pairs. We thus further extend OT by incorporating the deconvolution result, which, as mentioned above, maps a group of cells to a group of spots. As a result, scDOT integrates the two mentioned data modalities, single-cell and spatial transcriptomics, by simultaneously solving the deconvolution and OT problems. Specifically, given paired data modalities *X* and *Y*, representing gene expression profiles of a scRNA-seq and a spatial transcriptomics data respectively, scDOT simultaneously solves the deconvolution problem of estimating cell type fractions, *P*, of *c* cell types across *m* spots, and the spatial reconstruction problem of mapping *n* cells to their *m* spatial origins, resulting in a coupling matrix *γ*. These two solutions are constrained by the relation *γ* × *C* = *P*, where *C* is a binary matrix representing the cell type of each cell, encoded as a one-hot vector of size 1 × *c* across the total *n* cells. The two results, *P* and *γ*, are computed simultaneously in an iterative manner in order to improve each other’s accuracy. The problem is then formed as a bi-level optimization where the deconvolution and the spatial reconstruction are two inner optimization problems nested inside the outer optmimization that reflects the relation *γ* × *C* = *P*. See Supporting Methods for more details.

### 4.5 Inference of cell-cell spatial neighborhood graph

Utilizing the coupling matrix learned from optimal transport, we employed manifold alignment [47, 36] to project the single-cell data *X* and spatial coordinates *Z* ∈ ℝ^*m*×2^ of spatial transcriptomics data onto a common nonlinear subspace. This subspace preserves the correspondence between cells and spots, as well as the intrinsic similarity within each dataset. Consequently, in the common subspace, cells are represented in terms of both gene expression and spatial coordinates. Subsequently, we constructed a k-nearest neighbor graph (k-NNG) based on this new representation, which consists of the new coordinates in the common subspace for each cell. This allowed us to obtain the cell-cell spatial neighborhood graph. (In our experiments, we set *k* = 10.) The projections *f* and *g* resulting from manifold alignment serve as minimizers of the following optimization problem, which can be formulated as a generalized eigenvalue problem:

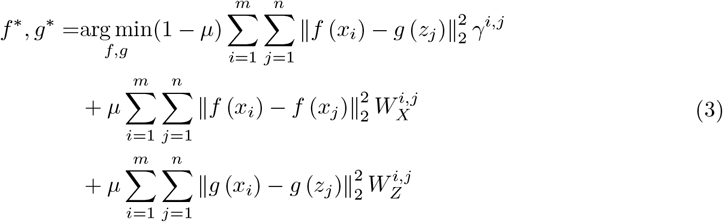

where *W*_*X*_ and *W*_*Z*_ are adjacent matrices of kNN graphs of *X* and *Z*, respectively.

## Notes

### Competing Interest Statement

The authors have declared no competing interest.

