## Supplementary Materials for "Optimal transport for mapping senescent cells in spatial transcriptomics"

[2]

[3]

[4]Machine Learning Department, School of Computer Science, Carnegie Mellon University, Pittsburgh, PA 15213, USA

#### A Supplementary Methods

##### A.1 Synthetic Data sets

**Synthetic data set 1** We first defined the covariance matrix for the grids. For two spots $s$ and $s'$, the covariance is defined as $[K]_{ss'} = \exp\left\{-\frac{\|t_s - t'_s\|_2^2}{\lambda\sigma^2}\right\}$, where $\sigma$ is the median value of spatial distance of all spot pairs, $\lambda$ is the similarity to adjust the smoothness of GP and is set to 0.1 by default. Next, we use $K$ as the kernel matrix of a GP to generate $c$ (number of cell types) samples for all cell types, simulating the cell type proportion at each spot $s$ as an "energy":

$$\phi^c \sim \mathcal{N}(\mathbf{0}, K) \tag{1}$$

Each $\phi^c$ is a cell-type energy vector aligned with spots. The proportion $\pi_s^c$ for each of the spot and each of the cell type is then driven by the energy,

$$\pi_s^c = \frac{\rho_c \exp\left\{\frac{\phi_s^c}{T}\right\}}{\sum_{c'} \rho_{c'} \exp\left\{\frac{\phi_s'}{T}\right\}}, \tag{2}$$

where T is a temperature parameter with a default value 1, $\rho_c$ represents the prior cell type distribution observed in the source data. A small value of $T$ tends to preserve the dominant cell type with the highest energy, while a large value of $T$ maintains the original cell type proportions. To ensure consistency with the sequencing method, each spot contains between 6 to 11 cells, and the total number of cells remains unchanged. To achieve this, we randomly assign one cell from the source data to a spot $s$, following the simulated cell proportion $\pi_s^c$. The mixed expression of each spot is then calculated as the mean expression of all cells within that spot.

**Synthetic data set 2** We followed previous studies [2, 3] and employed MERFISH to map the distribution of 135 specific genes in the MPOA brain tissue. Subsequent analysis using transcriptional clustering techniques on the gene expression measurements at the single-cell resolution revealed the presence of 9 major cell types. For simulating multi-cellular pixel-resolution spatial transcriptomics data, we aggregated the single-cell MERFISH data into pixels with an area of 100 μm$^2$. The simulation resulted in a paired single-cell and spatial transcriptomics data, containing 49142 cells across 135 genes and 3072 spots across 135 genes (consisting of 16 cells per spots on average). The ground truth also includes the annotations of 9 cell types and the spatial mapping of 49142 cells into 3072 spots.

#### A.2 Combination of deconvolution and mapping

The integration of cell type deconvolution and spatial mapping can be formulated as a bi-level optimization problem. This problem consists of two inner optimization problems: the deconvolution and the spatial reconstruction, which are nested inside the outer optimization problem representing the relationship between $\gamma \times C$ and $P$. The deconvolution problem is formulated as a NNLS optimization, while the spatial reconstruction problem is formulated as an OT problem:

$$
\begin{aligned}
\min_{\gamma^*, P^*} \quad & \|\gamma^* C - P^*\|_F \\
\text{s.t.} \quad & P^* \in \underset{P \in \mathbb{R}_+^{m \times c}}{\arg\min} \|Y - PS\|_F \\
& \gamma^* \in \underset{\gamma \in \mathbb{R}_+^{m \times n}}{\arg\min} \sum_{i,j} \gamma_{i,j} M_{i,j} + \lambda \Omega(\gamma) \\
& \qquad \text{s.t.}\, \gamma 1 = m; \gamma^T 1 = n; \gamma \geq 0
\end{aligned}
\tag{3}
$$

This bilevel optimization can be rendered differentiable, enabling efficient computation of backpropagation using the implicit differentiation theorem. It can be implemented by training a declarative neural network [1] in an end-to-end manner, allowing for iterative learning of the deconvolution and spatial reconstruction results until convergence is reached.

### B Supplementary Analysis

The cell-cell spatial neighborhood graph, denoted as $G$ (Figure 5b, 6c), is constructed from the cells of the single-cell data after being projected onto the common latent manifold using the manifold alignment technique (Methods), which preserves both the expression similarity and the spatial adjacency between cells. Consequently, an edge $uv$ exists in $G$ if either (1) cell $u$ and cell $v$ exhibit similar gene expression values, or (2) cells $u$ and $v$ are located in the same or adjacent spots, or (3) both (1) and (2) hold true. Although there are cases where cells with similar expression end up in the same or adjacent spots, this is not universally observed. To determine if the edges in $G$ are solely derived from gene expression similarity, we compare $G$ with another graph $G'$, which represents the cell-cell similarity graph based on similar expression only. Both $G$ and $G'$ are kNN Graphs with $k = 10$. Specifically, $G'$ is constructed from the original single-cell expression data $X$ as $G' = kNNG(X)$, where each cell is connected to its 10 most similar cells in terms of expression values. On the other hand, $G$ is constructed from the projection of cells into the latent manifold as $G = kNNG(f(X))$.

**For the paired dataset of an upper lobe slice:** In $G'$, there are $n = 134,610$ edges, while $G$ has $m = 80,086$ edges. There are only $p = 1,510$ overlapping edges between $G$ and $G'$. This indicates that only $p/m = 1.9\%$ of the edges in $G$ are attributed to expression similarity. The Jaccard index between $G'$ and $G$, calculated as $J = p/(n+m-p)$, is 0.007, indicating a substantial difference between the two graphs.

**For the unpaired dataset:** Using a similar calculation, we obtained the number of edges in $G$ and $G'$ as 290,160 and 502,722, respectively. There are 2,312 overlapping edges between the two graphs, which accounts for only 0.8% of the edges in $G$ resulting from expression similarity. The Jaccard index between the two graphs is 0.003, further highlighting a substantial difference between them.

### C Supplementary Figures

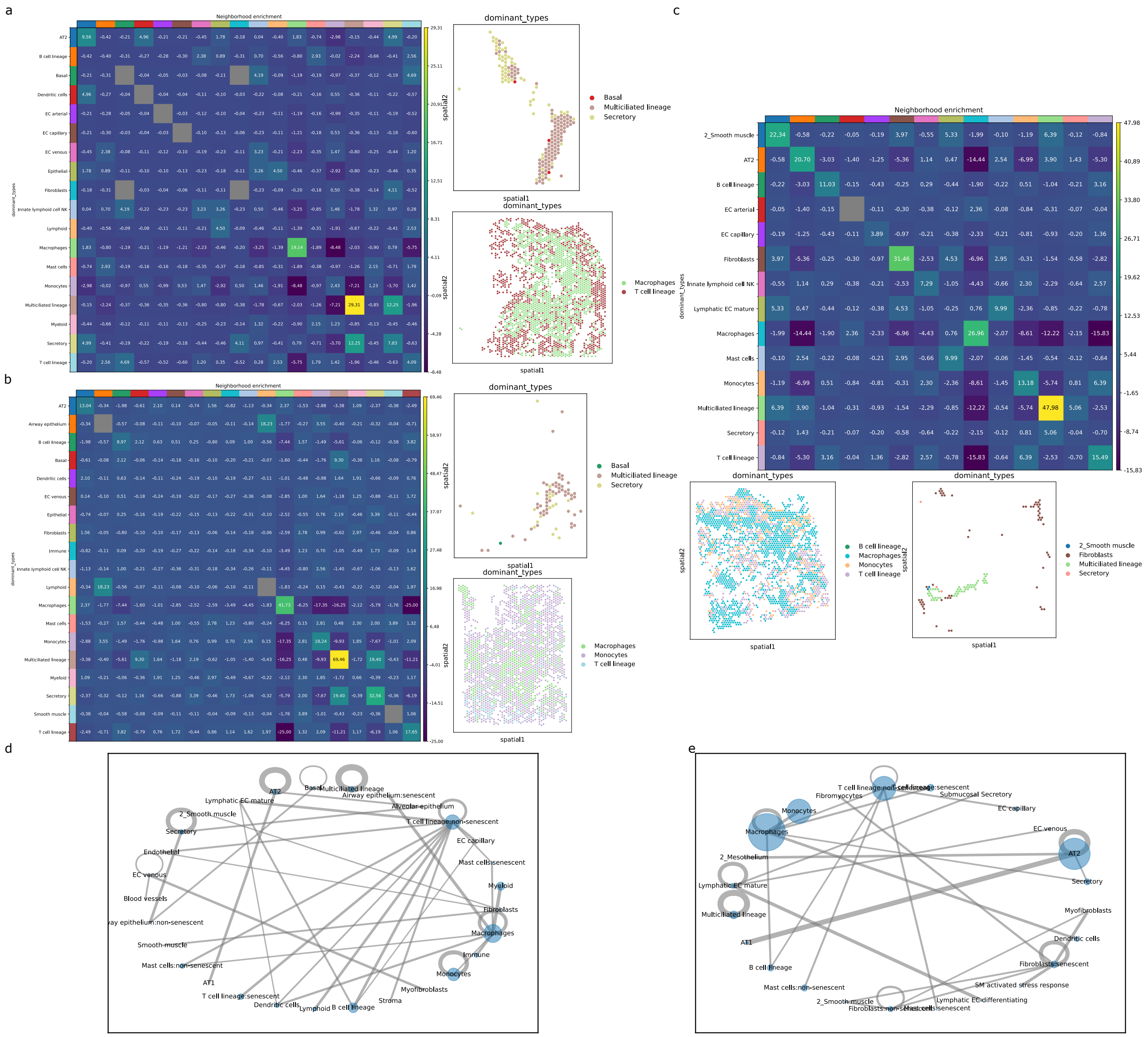


Supplementary Figure 1: Enrichment scores and cell-level neighboring counts are used to illustrate the spatial distribution patterns of cell types and their physical proximity.

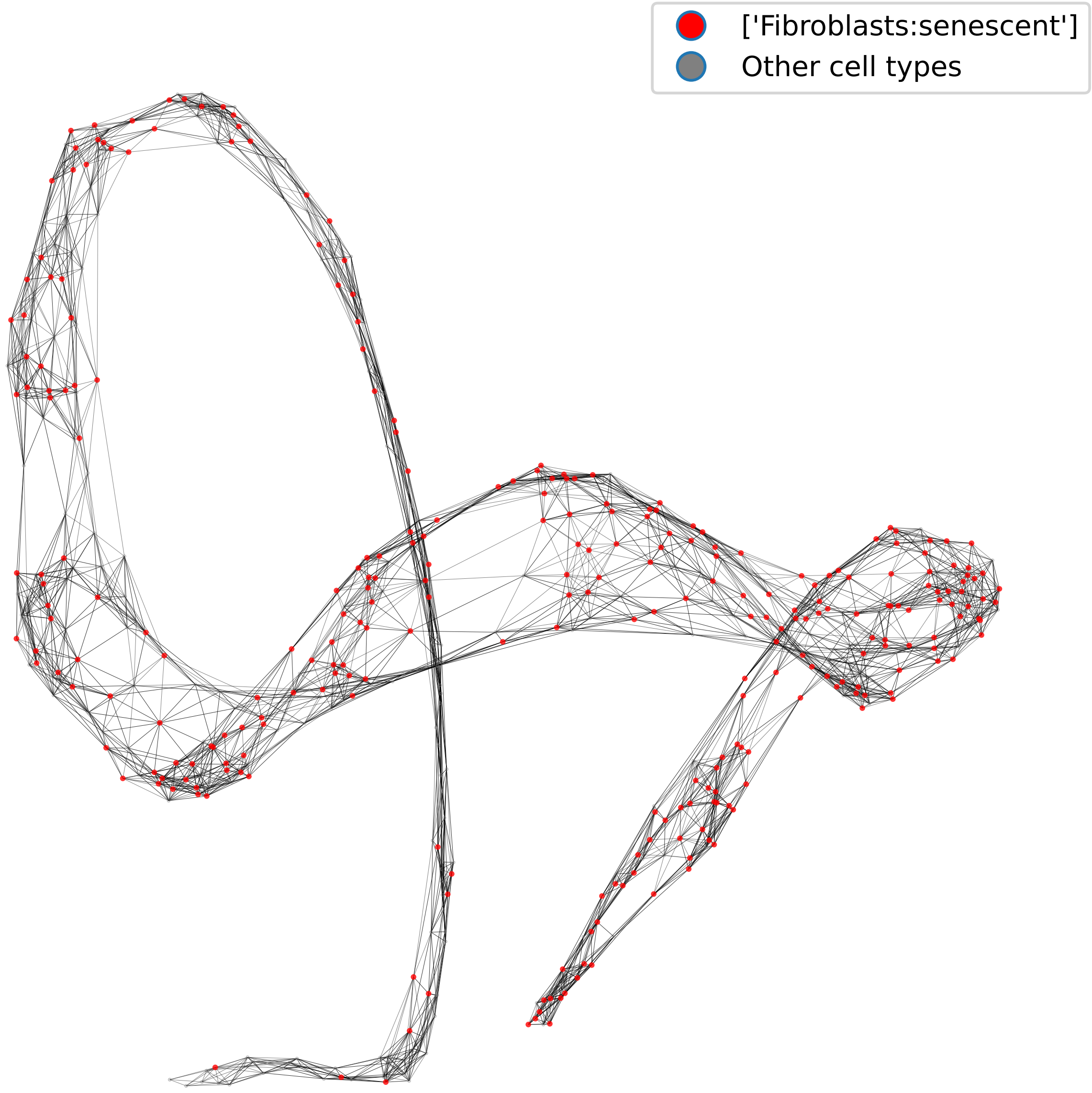


Supplementary Figure 2: Cell-to-cell neighborhood graph of senescent fibroblasts cells.

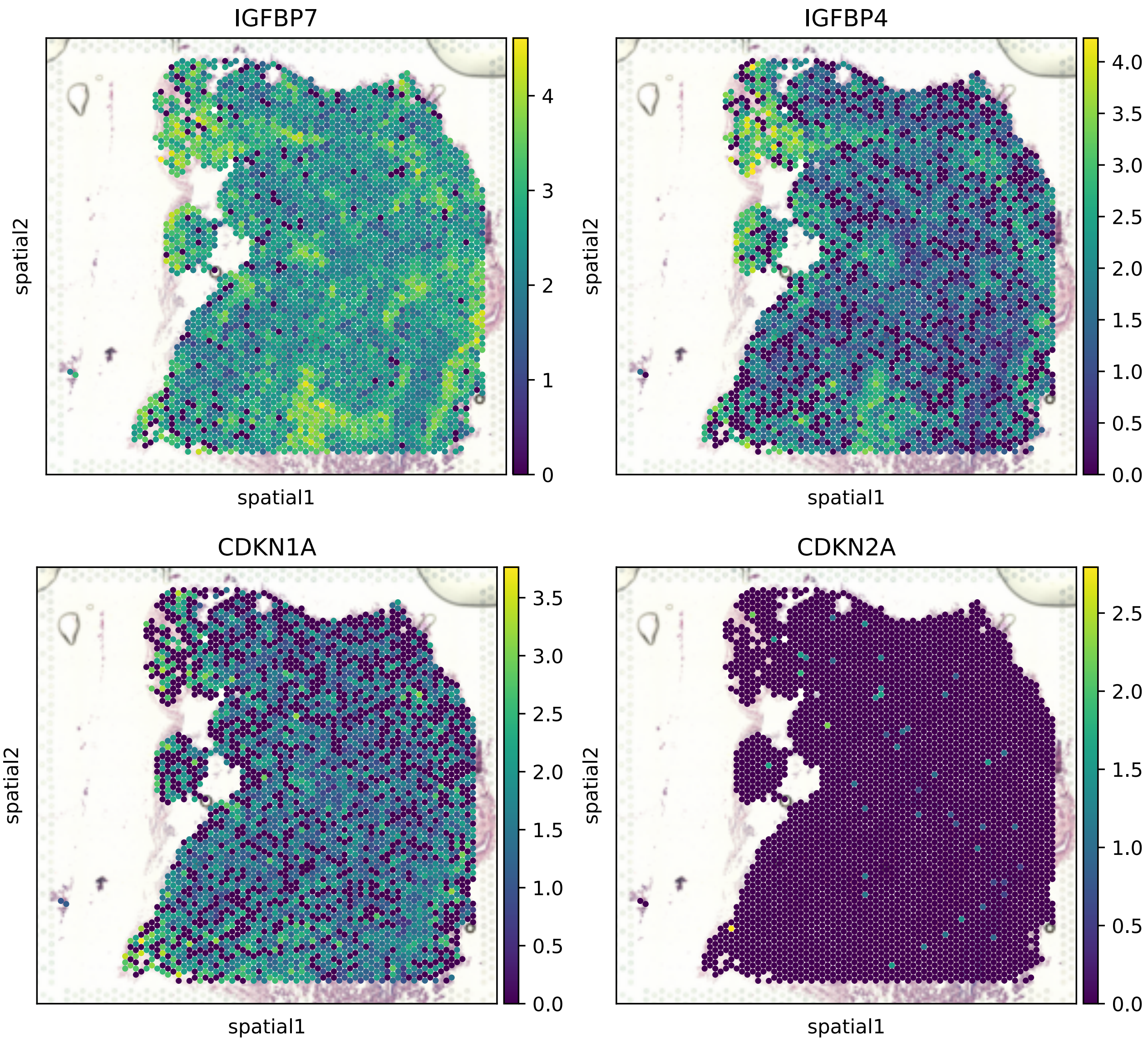


Supplementary Figure 3: Spatial distribution of selected senescent marker genes. IGFBP7 and IGFBP4 are among the 68 markers used for senescent reannotation. Although CDKN1A and CDKN2A are not included in this list, CDKN1A still exhibits high expression in the manually annotated region.
